# A Cross-Sectional Study of the Prevalence and Risk Factors of Soil Transmitted Helminthes Infection and Stunting Among School-aged Children in Ibadan

**DOI:** 10.1101/2020.01.16.908848

**Authors:** A. G. Ibrahim, M.K. Tijani, R. I. Nwuba

## Abstract

**Background:** In developing countries, infections caused by soil-transmitted helminthes (STH), such as *Ascaris, Trichuris* and hookworm, pose major public health problems among the school-age children, resulting in impaired physical growth such as stunting and thinness, and cognitive development. The aim of this study is to determine the prevalence of STH infections, stunting and thinness, and risk factors among school-age-children in Ibadan. This becomes highly imperative in order to serve as a guide on the prevention and control.

**Method:** A cross-sectional study was carried out in 8 primary schools at Ibadan, Oyo State Nigeria, between May and November 2018. All the school-age-children between the ages 5 and 18 years old (mean 10.4 ± 1.7 years), from primary one to six took part in the study. Demographic data were obtained and STH infections was analysed in single-stool samples by Kato-Katz. Anthropometric parameters were taken to calculate Height for-age Z score (HAZ) and Body-Mass-Index (BMI) for-age Z score (BAZ), in order to determine among school-aged-children stunting and thinness respectively.

**Results:** In overall, 458 school-age-children partook in the study. The prevalence of STH was 9.0%, out of which 7.6%, 2.8% and 1.5% were recorded for *Ascaris, Trichuris* and multiparasitism respectively. The overall prevalence obtained revealed that the results of stunting and thinness (HAZ < -2SD, BAZ < -2SD) were 24.7% and 27.3% respectively based on the WHO Child Growth Standards (2007). Notably, ages of the children (*P*< 0.01), their classes (*P*=0.05), different schools (P=0.003), washing of hands after toileting (*P* = 0.05) were important risk factors determining STH infection, HAZ and BAZ.

**Conclusion:** The study showed that specific risk factors among school children in the studied area will make them vulnerable with high risk of STH infection, HAZ and BAZ. Effective prevention and control strategies can be well planned when risk factors and dynamics of transmission in vulnerable groups have been painstakingly identified.

**Summary:** *Ascaris, Trichuris* and hookworm are responsible for major public health problems among the school-age children (SAC); this has led to stunting and thinness, and impaired cognitive development. With the aim to determine prevalence of STH infections, stunting, thinness and associated risk factors, a cross-sectional study of STH infection in 8 primary schools at Ibadan, Oyo State Nigeria was conducted in 2018. School-aged children between the ages of 5 and 18 were enrolled, demographic data, stool samples and anthropometric parameters were obtained in order to determine STH infection and nutritional status. 9.0% was the prevalence of STH, the prevalence of 7.6%, 2.8% and 1.5% were recorded for *Ascaris, Trichuris* and multiparasitism respectively, while 24.7% and 27.3% were obtained for stunting and thinness respectively. The children’s age, hand washing after toileting and locations of the different schools were significantly correlated with STH infection, HAZ and BAZ. This study is highly imperative because its shows some risk factors associated with STH infection, HAZ and BAZ among SAC, this can serve as a guide on the prevention and control among SAC.

## INTRODUCTION

Among the most common infections worldwide which affect the poorest and most deprived communities was the soil-transmitted helminthes (STH) infections, the most prevalent neglected tropical diseases [16]. Roundworm (*Ascaris lumbricoides*), hookworms (*Necator americanus* and *Ancylostoma duodenale*) and whipworm (*Trichuris trichiura*) are the main species that infect people. In areas where there is poor hygiene, STH transmission is by ingesting eggs present in human faeces which invariably contaminate the soil, or through skin penetration. World Health Organization [25] estimation indicated more than 5 billion people are at risk and over 2 billion people in the world are infected with these parasites [3]. School Aged Children (SAC) of about 1 billion were at risk of being infected with at least one STH species [15]. There is the concentration of STH infections in impoverished regions such as the tropical and sub-tropical, where clean water and provision of sanitation is deficient and the practices of poor personal hygiene are common [27,4,20]. Prominent symptoms of infected individuals with STH are anaemia, stunting growth, pain in the abdominal, diarrhoea, intellectual retardation, and death when infections are chronic and untreated [2,7,26]. SAC living in an endemic environment where sanitation is poor are mostly at risk of STHs infection and this infections has immensely contributed to early childhood debilitation [18,30]. A common health problem in the world is malnourishment; one out of three people are malnourished [29]. STH infections are found to predispose one to malnutrition; For instance, increase in the susceptibility of SAC to parasitic infection is due to their poor intake of nutrients leading to their poor growth and development [14,29]. When STHs feed on host tissue, it will lead to deficiency of protein and iron, loss of appetite, diarrhoea, dysentery and increment in malabsorption, thereby leading to malnutrition [28]. Stunting or low Height-for-Age Z score (HAZ) and low Body Mass Index (BMI)-for-age Z score (BAZ), are good indicators of malnutrition which can be used to represent the status of stressed chronic nutritional [18]. In some studies, different countries have reported that STH such as *Ascaris, Trichuris* and multi-parasitism were linked with HAZ in school-age-children [1,17]. The aim of the study was to determine the prevalence of STH infections, stunting and thinness and their associated risk factors among SAC in Ibadan. This becomes highly imperative in order to provide guide on how to prevent and control STH among SAC in Ibadan.

## MATERIALS AND METHODS

### Study Area

This was a cross-sectional study conducted among school-aged children in Ibadan, Oyo State between May and November 2018. The city has a population of about 3.8 million people. It has 11 Local Government Area, out of which 5 are urban and 6 are semi-urban. The principal inhabitants are the Yorubas.

### Study population

School-age children aged between 5 and 18 years old (mean age 10.4 ± 1.7 years), from primary one to six from four Local Government Areas (LGA), namely: Ibadan South-East LGA and Ibadan North-West LGA (urban LGA), Akinyele LGA and Ido LGA (semi-urban LGA), took part in the study. Two schools and a minimum of 50 pupils in each school were randomly selected per LGA: IBSE-SCH1 and IBSE-SCH2; IBNW-SCH1 and IBNW-SCH2; AKIN-SCH1 and AKIN-SCH2; IDO-SCH1 and IDO-SCH2.

#### Sample size determination

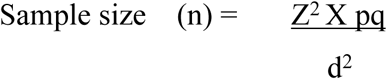

where Z=1.96, p=0.359, q=0.641, d= 0.05

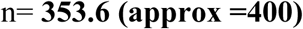

A population of randomly selected 458 school-aged children eventually took part in the study.

### Ethical Approval and Questionnaires for collection of demographic history

Approvals for the protocol were obtained from the UI/UCH Ethical Review committee, College of Medicine, University of Ibadan (NHREC/05/01/2008a); Oyo State Ministry of Health (AD 13/479/614), Oyo State Ministry of Education (EDU. 215T2/3) and Oyo State Universal Primary Education Board (SUPEB) (SUBEB/G.1157^T^/32). Parent Teacher Association meetings were held before the commencement of the study. Written informed consents and assents were obtained from the parents and their children/ward respectively.

The inclusion criteria:

i. The individual will be willing and their parent or guardians will give consent

The exclusion criteria:

i. Children whose parents or guardians failed to give their consent to the study
ii. Children who were on drugs for any infection

All individuals in the study were interviewed with a structured questionnaire. Demographic information for age, sex, education, past history of worm infection and treatment, socio-economic status of the parent, mother’s level of education, frequency of treatment, availability of toilet facilities, hand washing practices etc were obtained with the questionnaire.

### Collection and examination of stool samples

Children who met the inclusion criteria were given universal bottles (100ml) and were instructed on how to do proper faecal collection. Helminth eggs in stool were quantitatively estimated using the Kato-Katz thick smear technique. The intensity of STH infections was expressed as the number of eggs per gram (epg) of faeces: *A. lumbricoides*, 1–4,999 epg, 5,000–49,999 epg and ≥50,000 epg for light, moderate and heavy respectively; while in *T. trichiura* infection the presence of 0-999 epg, 1,000-10,000 epg and >10,000 epg for light, moderate and heavy respectively [10]. All children positive for STH infection were treated with benzimidazole (mebendazole) by a health officer.

### Assessment of Anthropometric among SAC

The children’s height and weight were measured; age and sex were collected via questionnaire to determine their anthropometric indices. Heights of the children were measured to 0.1 centimeters (cm) using a height measuring tape while the pupils’ weights were recorded using a scale to the nearest 0.1 kilograms (kg) using a weighing scale. As indicators of under-nutrition using the WHO Anthroplus software, HAZ and BAZ were estimated to determine underweight and stunting status of the SAC respectively. These calculations will take children’s weight and height into account [23]. HAZ and BAZ scores were classified as normal nutrition (≥ −1.01 to −2.00 Standard Deviation [SD]) and under-nutrition (< −2.01 SD) [6,11].

### Data analysis

Microsoft excel and IBM SPSS version 20.0 were used for statistical analyses, using p<0.05 as statistical significance. Descriptive statistics was performed selecting variables for consideration in univariate and multivariate regression models, STH status as dependent variable. World Health Organization Child Growth Standards was used to analyze Anthropometric data (22). Analysis of prevalence and intensity of STH was done using Univariate and multi-variate analyses with the socio-demographic indicators of age, sex, risk-behaviors such as hand-washing, previous history and frequency of deworming treatment, and use of toilets etc.

## RESULTS

### Characteristics of the SAC

Out of the four Local Government Areas that was studied, 458 eligible children from 8 schools provided fresh stool samples. The characteristic of the studied population is represented in Table 1. 10.4 years (range 5–18) was the mean age, and boys were slightly more than girls (51.2% vs 48.8%). Access to sewer system at homes (96.3%) and at schools (85.8%) was present for the children evaluated. Majority of the children (81.4%) had been treated with an anthelminthic drug in the past 12 months.

**Table 1:**
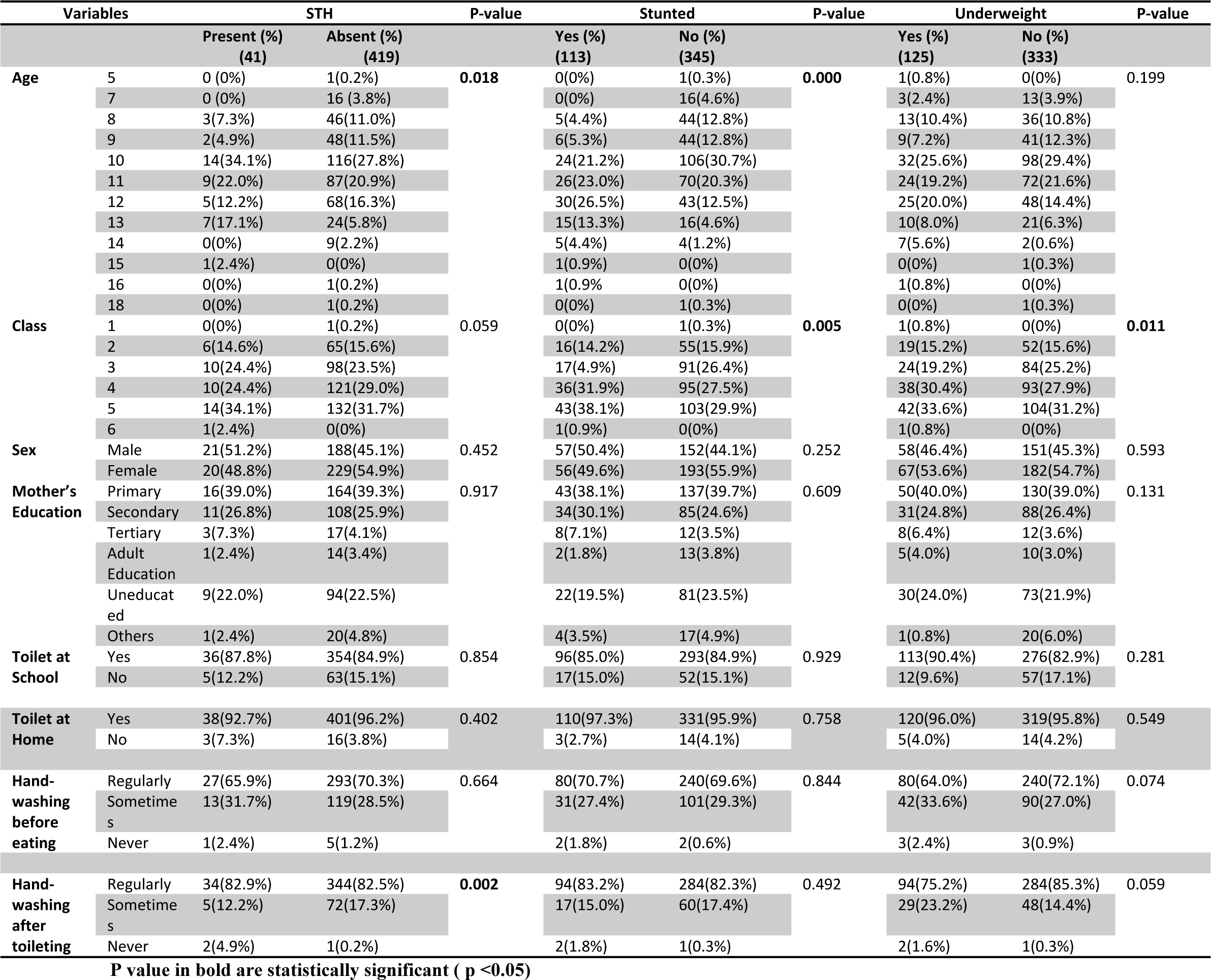
Socio demographic characteristics, soil transmitted helminth and nutritional status of school-age-children in eight primary schools in Ibadan

### Prevalence and intensity of Soil Transmitted Helminthiasis

The prevalence of soil transmitted helminthiasis with at least one parasites was 9.0% (95% CI 6.6 to 11.6). Most frequent infections were with *A. lumbricoides* 7.6% (95% CI 5.2 to 10.0) followed by *T. trichiura* (2.8%, 95% CI 1.5 to 4.6) and 1.5% (95% CI 5.47 to 55.18) for multiparasitism, hookworm was not detected in this study. Parasite intensity for ascariasis and trichuriasis were determined using direct egg per gram count (epg) according WHO, 2002. In this study, 18 (3.9%) and 17 (3.7%) participants had light and moderate *A. lumbricoides* infection respectively, while for *T. trichiura* infection, 11 (2.4%) and 2 (0.4%) participants had light and moderate infections respectively. In this study, no record of heavy infection was found for the two STHs. Mean intensity of infections were 6202 eggs/g (epg) with a range of 24–36000 and 436 egg/gram (epg) (range: 24-3696) for *A. lumbricoides* and *T. trichiura* respectively (24).

### Prevalence and intensity of STH infections by selected schools

Of the 8 selected schools in the four Local Government in Ibadan, four schools were located in the urban region (IBSE-SCH1 and IBSE-SCH2; IBNW-SCH1 and IBNW-SCH2) while the other four were in the semi-urban region (AKIN-SCH1 and AKIN-SCH2; IDO-SCH1 and IDO-SCH2). The prevalence of STH infections observed between the studied eight schools was significant (p<0.000) (Fig. 1). In this study, IBSE-SCH1 was observed to have the highest rates of ascariasis and trichuriasis (15%, and 10%, respectively) (Fig. 2) while in other schools varying rates between 0% and 8% prevalence was observed for each of the two parasites. Among infected children, light and moderate infection intensities with *A. lumbricoides* (3.9% and 3.7%) and *T. trichiura* (2.4% and 0.4%) were detected (Fig. 3). In this study, there was no record of heavy infection intensities with *A. lumbricoides*, and *T. trichiura* in the all the schools visited.

**Fig. 1:**
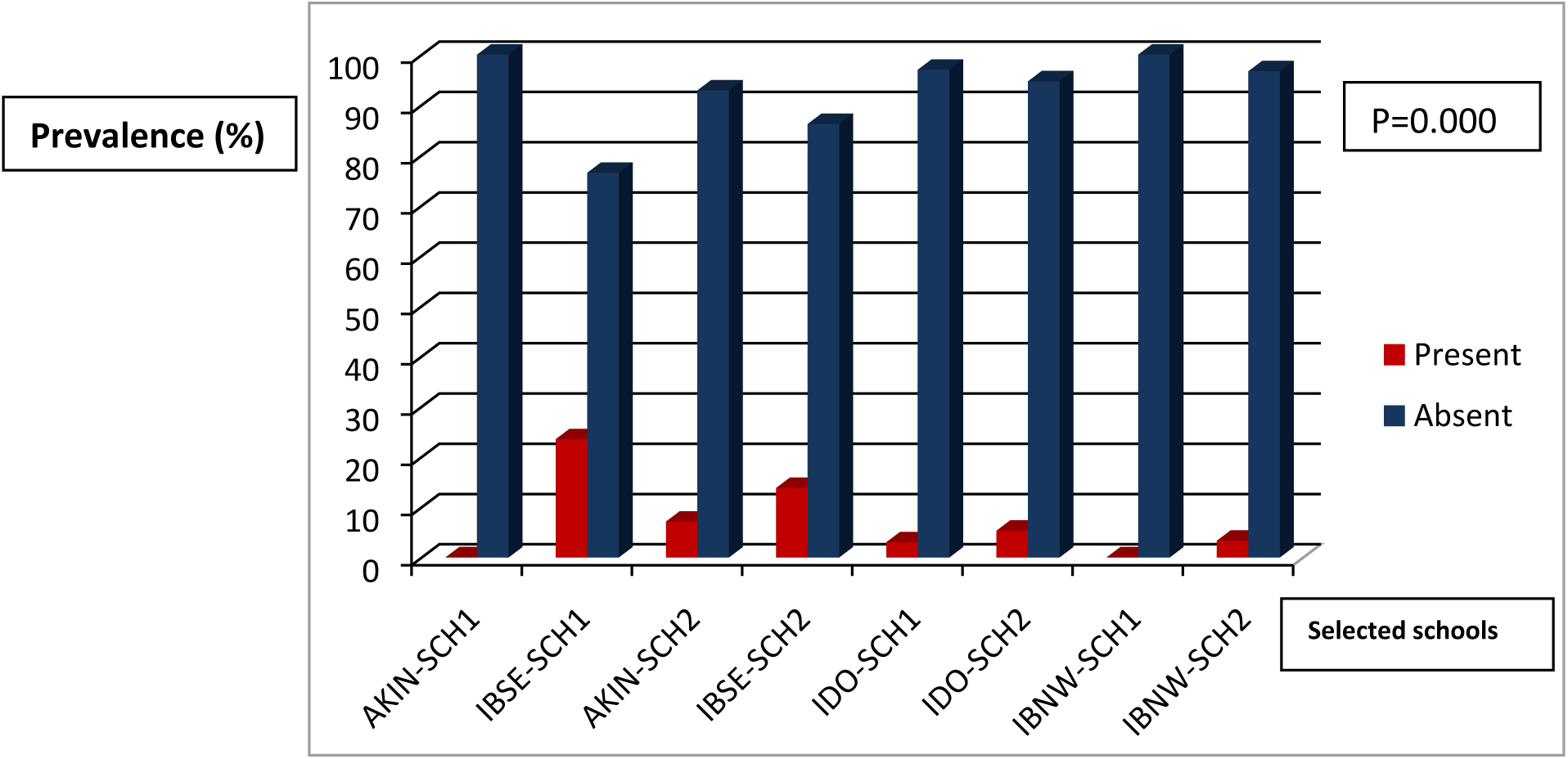
Prevalence of STH infections by selected schools.

**Fig. 2:**
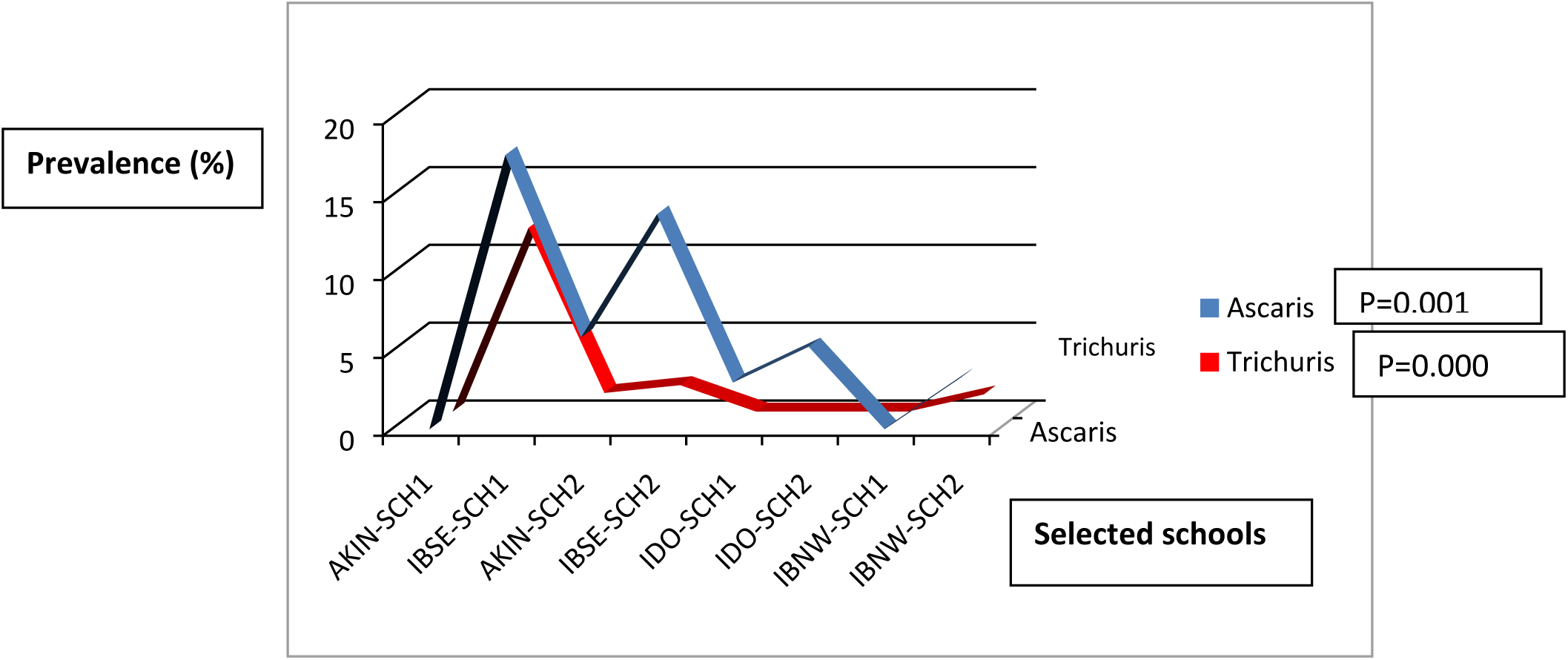
Prevalence of *Ascaris* and *Trichuris* infections by selected schools.

**Figure 3:**
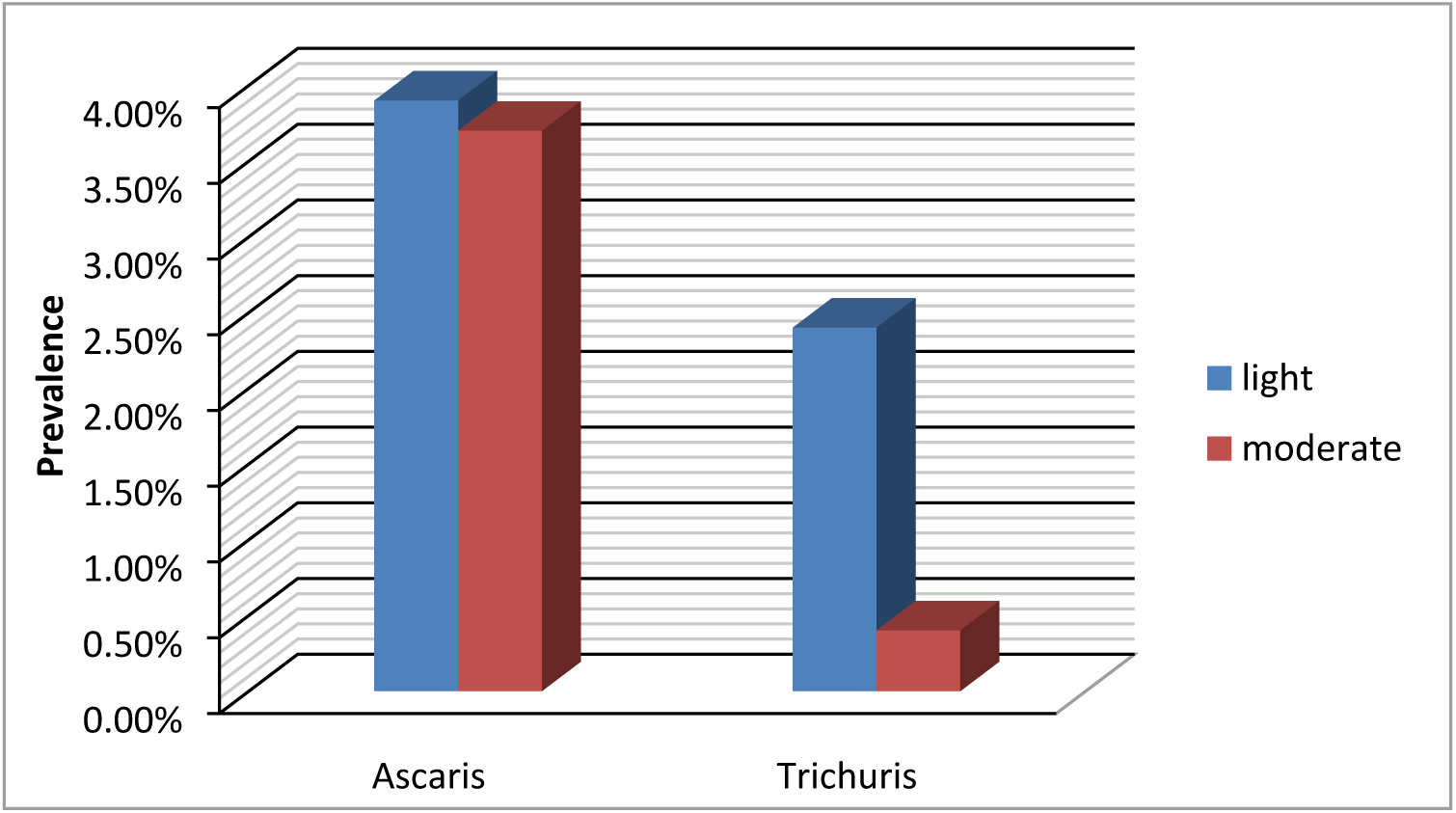
Intensity of *Ascaris* and *Trichuris* among the SAC in Ibadan. N.B: The intensity of both infections are generally higher among males. For instance, in Ascaris, light: male=28.6%, female=22.9%; moderate: male=25.7%, female=22.9%; in Trichuris males=46.2%, females=38.5% and males=15.4%, females=0% for light and moderate respectively). There is no significant difference. The intensity is higher in ages 8-11 years for both infection compared to other ages

The prevalence and intensity patterns for *Ascaris* and *Trichuris* vary by age of the SAC in this study, children between the ages 8–11 years had high prevalence of *Ascaris* and *Trichuris* but was lower in older children; this was not statistically significant (Fig. 3).

### Prevalence of malnutrition

In this study, the prevalence of HAZ (<-2 SD) and BAZ (<-2 SD) was 24.7% (95% CI 94.5 to 98) and 27.3% (95% CI 29.5 to 38.6), respectively. The prevalence of stunting and thinness varied significantly, p= 0.003, p= 0.013, and IBSE-SCH2 was observed to have the highest prevalence of malnutrition, (23% HAZ) and (20% BAZ) respectively (Fig. 4).

**Figure 4:**
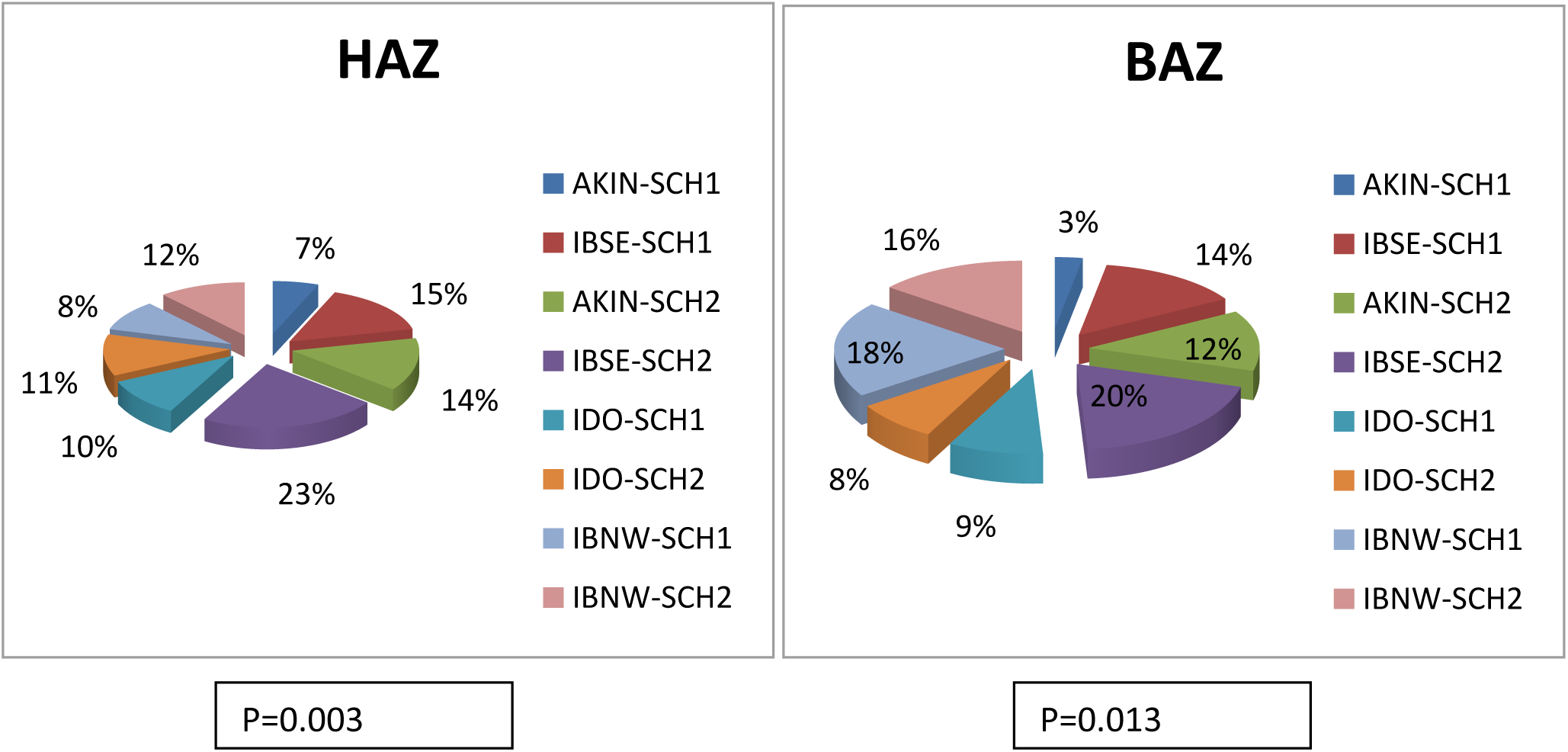
Prevalence of HAZ in the selected schools. Prevalence of BAZ in the selected schools

### Relationship between STH infections and malnutrition

In this study, it was observed that the presence of trichuriasis among school aged children had negative impact on their weights and heights but this was not statistically significant (p=0.283 and p=0.063 respectively). Also, ascariasis had a significant (p=0.005) negative impact on their heights (Tab. 2).

**Table 2:**
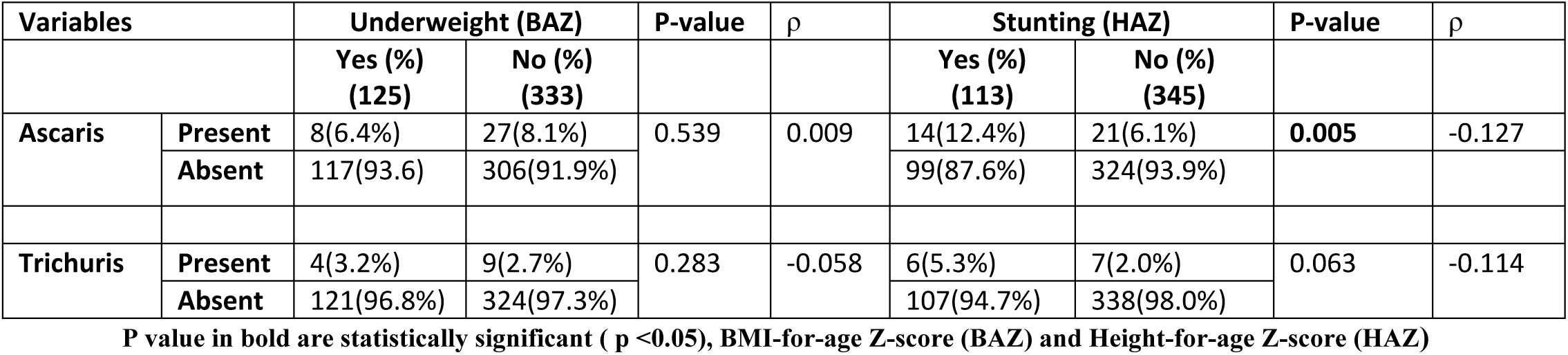
Association between STH & BAZ, HAZ.

| Variables |  | Underweight (BAZ) |  | P-value | ρ | Stunting (HAZ) |  | P-value | ρ |
| --- | --- | --- | --- | --- | --- | --- | --- | --- | --- |
|  |  | Yes (%)<br>(125) | No (%)<br>(333) |  |  | Yes (%)<br>(113) | No (%)<br>(345) |  |  |
| Ascaris | Present | 8(6.4%) | 27(8.1%) | 0.539 | 0.009 | 14(12.4%) | 21(6.1%) | **0.005** | -0.127 |
|  | Absent | 117(93.6) | 306(91.9%) |  |  | 99(87.6%) | 324(93.9%) |  |  |
| Trichuris | Present | 4(3.2%) | 9(2.7%) | 0.283 | -0.058 | 6(5.3%) | 7(2.0%) | 0.063 | -0.114 |
|  | Absent | 121(96.8%) | 324(97.3%) |  |  | 107(94.7%) | 338(98.0%) |  |  |

P value in bold are statistically significant ( p <0.05), BMI-for-age Z-score (BAZ) and Height-for-age Z-score (HAZ)

### Relationship between STH & BAZ, HAZ

The result of univariate and multivariate analysis is represented in Table 3. Associated Risk factors for HAZ and BAZ were: Age (OR = 1.705; 95%CI: 1.426 - 2.039); class (OR = 0.713; 95%CI: 0.547 – 0.930); access to toilet at home (OR = 0.425; 95%CI: 0.206 – 0.875); washing of hands after using the toilet (OR = 2.122; 95%CI: 1.298 – 3.467).

**Table 3:** Univariate and multivariate logistic regression of factors associated with HAZ and BAZ among school aged children in Ibadan

| Variables | HAZ (n=454)<br>Coefficient (β) (95% CI) | P-value | BAZ (n=454)<br>Coefficient (β) (95% CI) | P-value |
| --- | --- | --- | --- | --- |
| Age | 1.705 (1.426-2.039) | **0.000** | 1.187 (1.018-1.383) | **0.029** |
| Class | 0.713 (0.547-0.930) | **0.012** | 0.868 (0.680-1.107) | 0.253 |
| Sex | 0.897 (0.567-0.930) | 0.642 | 1.007 (0.657-1.544) | 0.974 |
| Toilet at school | 0.821 (0.407-1.655) | 0.581 | 0.755 (0.353-1.617) | 0.470 |
| Toilet at home | 0.818 (0.448-1.494) | 0.514 | 0.425 (0.206-0.875) | **0.020** |
| Hand-washing after toileting | 1.120 (0.643-1.951) | 0.689 | 2.122 (1.298-3.467) | **0.003** |

P value in bold are statistically significant ( p <0.05)

## Discussion

This study recorded a prevalence of 41 (9.0%) for helminth infections, out of which *A. lumbricoides* (7.6%) was the highest while *T. trichiura* was 2.8%. This is in contrast to earlier studies conducted in Igbo-Ora, Oyo State Nigeria by Ibrahim *et al*., (2015) [8] who recorded hookworm as the highest STH, but in line with by Ojurongbe *et al*., (2011) [13]. More so, the low prevalence of STH, 9%, recorded in this study is in contrast to Ikeoluwa *et al*., (2015) [9] who recorded 35.9%, this cannot be far fetch from the recent mass drug administration that recently took place all over the state around four months before the commencement of this study.

In this study, the prevalence of STH infection was found to have significant differences in the different schools sampled. Ibadan South-East was observed to have the highest prevalence of the STH infection; for instance, 23.5% and 13.8% prevalence was observed in IBSE-SCH1 and IBSE-SCH2 respectively. In collaboration with another study [9], Ibadan South-East can be tagged as a hotspot for STH infection. This may be as results of the schools were located in a densely populated area and also the unhygienic lifestyles of both the pupils and their parents which enhance the spread of the infection.

In this study, prevalence of STH across the ages was significant, ages 8-11 years was dominant. This age group is known to be active and may be engaged in activities, such as not washing of hands after toileting etc, which may promote the spread of the infection; More so, the males’ pupils (51.2 %) were insignificantly more than the females’ pupils (48.8%). This is corroborated by Ikeoluwa et al., (2015) [9] but in contrast to Moncayo *et al*., (2018) [12].

In this study, we recorded low prevalence of STH (9.0%), compared to HAZ and BAZ, 24.7% and 27.3% respectively, with IBNW-SCH2, BAZ=20% and HAZ=23% having the greatest prevalence in the sample. Unlike STH, each visited school has at least a case of malnutrition; prolong history of being malnourished or other infections which can predisposed them malnutrition may be responsible [18].

In this study, the presence of *Ascaris* leads to stunting among the infected children and this was significant, whereas *Trichuris* infections also negatively affect children’s growth and development leading to stunted growth and wasting, but this has no significant difference. STH could affect growth in SAC using several mechanisms which may include reduction in food intake, malabsorption and reduced appetite [5]. This is in accordance with previous studies which indicate a correlation between malnutrition and STH infection [5, 19].

In this study, hand-washing after toileting, ages of the children, their different classes and the accessibility of the children to toilet at home were important risk factors for the spread of ascariasis and trichuriasis which invariably affect the children’s growth and development. Thus, if teachers are better grounded/ educated on WASH and proper health and nutrition behaviours, they will definitely help in transmitting the knowledge to both the pupils and their parents; this will definitely aid in reducing and/or eradicating STH infections among SAC. Therefore, educational interventions on health and nutrition with teachers and SAC cannot be overemphasized.

In conclusion, the accurate and up-to-dated information on the prevalence data obtained in this study will aid in mapping out the current situation of STH infections among SAC in Ibadan. Furthermore, insight into the nutritional status of school-age-children living in semi-urban and urbanized metropolis of Ibadan was uniquely provided for in this study. Nonetheless, more researches into the health impact of STH-polyparasitism and other infections that can predispose SAC to malnutrition should be carried out. Hopefully, the findings presented here will be helpful in formulating current STH prevention and control in Ibadan among school age children.

## Supporting information

**S1 Checklist** STROBE statement.

(DOC)

## Acknowledgement

We are grateful to the Headmasters and Headmistress of the eight sampled-schools, their teachers, all the pupils who participated in the study and their parents for their enthusiastic participation.

